# Development of a *De Novo* Protein Binder that Inhibits the Alpha Kinase eEF2K

**DOI:** 10.1101/2024.12.10.627789

**Authors:** Kody A. Klupt, Ethan Belrose, Zongchao Jia

**Affiliations:** Department of Biomedical and Molecular Sciences, Queen’s University, Kingston, ON, K7L 3N6, Canada

**Keywords:** eEF2K, Alpha Kinase, De Novo Protein, Protein Engineering, Kinase Inhibitor, Calmodulin

## Abstract

Elongation factor 2 kinase (eEF2K) is calmodulin activated and phosphorylates eEF2, a GTPase, that regulates global translation. When eEF2K phosphorylates eEF2, protein translation is halted. This process may be critical to studying how diseases like cancer dysregulate protein synthesis. eEF2K is an alpha kinase and not targeted by conventional kinase inhibitors. Traditional methods of structure-based drug design are incredibly time consuming and expensive, which may involve screening large libraries of small molecules. We have generated *de novo* small binder proteins (∼10kDa) - using RFDiffusion and ProteinMPNN. One promising *de novo* binder protein we produced, ‘CAM1’ binds to a hydrophobic patch on the calmodulin binding domain of eEF2K with nanomolar affinity as determined by isothermal titration calorimetry. This binder, *in vitro*, significantly reduces eEF2 peptide phosphorylation, comparable to the gold-standard small molecule eEF2K inhibitor, A-484954. The predicted structure of CAM1 is a helical bundle which has been confirmed by circular dichroism spectroscopy. Impressively, CAM1 has a melting temperature >80^0^C, and is produced recombinantly in bacteria, greater than 5 mg / culture liter. We have also determined that CAM1 transfection significantly reduces mammalian HeLa cell proliferation comparable to A-484954 treatment and inhibits the phosphorylation of eEF2. Our *de novo* binder, the first to our knowledge to inhibit an alpha kinase, and the first non-competitive eEF2K inhibitor, establishes an alternative method of targeting atypical kinase activity.

## INTRODUCTION

Eukaryotic elongation factor 2 kinase (eEF2K) is a calmodulin (CaM) activated kinase that phosphorylates the threonine 56 residue of the GTPase eEF2^1–3^. eEF2 when phosphorylated, is unable to bind the ribosome and catalyze the translocation of the elongating peptide between the A and P ribosomal sites and global peptide translation is halted^4^ (Figure 1a). eEF2K is an alpha kinase, as such its sequence is dissimilar to the hundreds of conventional protein kinases, and resulting structural differences protect eEF2K from inhibition by most conventional kinase inhibitors^5–8^. eEF2K contains several domains, including an N-terminal CaM binding domain (CBD), alpha kinase domain, and C-terminal eEF2 binding domain^9,10^ (Figure S1a). The interaction between the CBD, CaM, and alpha kinase domain is critical for kinase activation, and has been extensively characterized both enzymatically and structurally. CaM binding to the CBD is calcium dependent, and calcium induced structural changes of CaM permit tighter binding to the CBD^1,11,12^. The CaM binding to eEF2K stabilizes its active kinase conformation and this structural shift is critical for activation. The proposed mechanism of eEF2K activation includes autophosphorylation, whereby, the CaM binding enhances the affinity for Thr348 of eEF2K to be rapidly phosphorylated, which then becomes associated with a basic residue pocket on the alpha kinase domain^13^. Thr348 is located on an important regulatory loop which is believed to be the downstream substrate of a variety of signaling pathways implicated in disease^14^.

**Figure 1.**
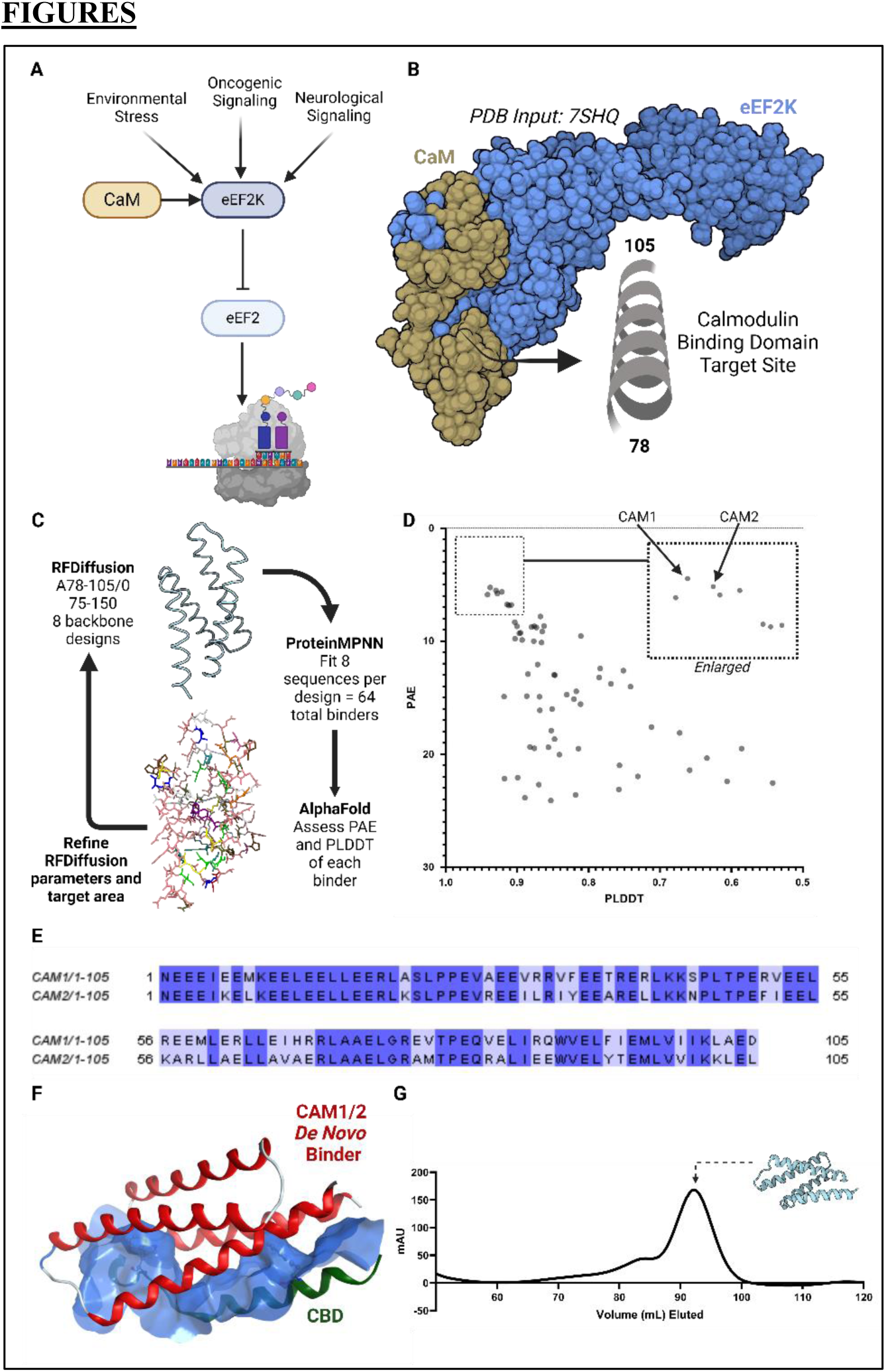
Design of a *de novo* protein binder against the CBD of eEF2K. eEF2K which phosphorylates eEF2 is A) a downstream target to a variety of signaling cascades implicated in diseases like cancer. B) Structural overview of eEF2K and CaM (PDB: 7SHQ). CaM interacts with the CBD of eEF2K, a helical region. C) Overview of *de novo* design and optimization process with RFDiffusion, ProteinMPNN, and AlphaFold. D) 64 binders with sequence-structure pairs were assessed with AlphaFold scores PAE and PLDDT. CAM1/2 have the lowest PAE scores with PLDDT above 0.9. E) Sequence alignment of CAM1 and CAM2. F) Predicted backbone structure of CAM1/2 (red) interacting with the CBD of eEF2K (green). Blue surface of CBD demonstrates an intimate association with CAM1/2. Tryptophan residues of CBD displayed. G) Representative size-exclusion chromatogram of CAM1 recombinant purification.

Inhibition of eEF2K has been extensively studied as a potential therapeutic strategy for various diseases, including cardiovascular conditions, neurological disorders, and cancer^14^. As the sole member of an atypical kinase family activated by CaM, targeting the CaM mediated activation mechanism of eEF2K could yield significant therapeutic benefits. eEF2K plays a critical role in cancer by regulating protein synthesis and enabling cancer cells to adapt to stress, such as nutrient deprivation and hypoxia^15^. By inhibiting energy-intensive peptide chain elongation, eEF2K can conserve cellular resources, and promotes glycolysis (Warburg effect), therefore supporting survival under adverse conditions^16,17^. Its activity has been linked to tumor progression, metastasis, and resistance to chemotherapy and radiotherapy. Elevated eEF2K expression is associated with poor patient survival in cancers like lung and breast cancer^18,19^.

Conventional kinase inhibitors which may be non-selective, are unable to inhibit eEF2K kinase activity, as demonstrated previously with staurosporine (IC>50 uM)^20^. An effort to inhibit eEF2K by small-molecule inhibitors which has been ongoing for several decades began with the discovery that rottlerin (IC50 = 5.3 uM) can restrict eEF2K activity^20^. The design of targeted therapeutics such as NH125 (IC50 = 60 nM) was successful *in vitro*, however, later work argued NH125 was responsible for the induction of eEF2 phosphorylation^21,22^. Development of A-484954, the existing gold standard small molecule inhibitor of eEF2K has an IC50 of 2.8 uM, inhibits eEF2 phosphorylation, and is commercially available as a research tool^22^. To improve A-484954 potency, researchers have modified this compound, for example, connecting the compound to an E3 ligand for targeted eEF2K degradation by the proteasome (PROTAC method), demonstrating both reduction in total eEF2K and eEF2 phosphorylation in breast cancer cells^23^. Despite these efforts, traditional methods of small molecule structure-based drug design nonetheless contain obstacles including the need for high throughput compound screening and at times, complex chemical synthesis or modification.

It is worthwhile exploring alternative strategies to inhibit the CaM mediated activation of eEF2K such as targeted *de novo* protein binder design. Compared to small molecules however, protein structure prediction is computationally more difficult due to the rich energetic landscape of protein folding^24^. Within the last several decades, a major challenge has been developing novel proteins with targeted functions, backbones, and sequences unrelated to existing proteins in the proteome. Such proteins that do not exist in nature are categorized as *de novo,* yet are still dictated by the biophysical principles governing protein folding^25^. *De novo* protein design has advanced significantly within the last few decades from designing novel folds to producing enzymes with enhanced catalytic activity^26,27^.

Popular programs used today for *de novo* binder design, which builds a specific binder protein onto a protein target, include RFDiffusion (RF for Rosetta Fold), and ProteinMPNN (MPNN for message passing neural network)^28,29^. RFDiffusion iteratively generates a protein by denoising protein structure data to create a diverse backbone output that reflects a protein-like structure but does not exist naturally. Side chains can then be fitted into the backbone with ProteinMPNN, generating a structure-sequence pair^28,29^. Together, such programs may produce a *de novo* binder that has an intimate fit with its target protein region and would be expected to have a good binding affinity.

In this paper, we apply *de novo* protein design to generate a binder that targets the CBD of eEF2K. Herein, we seek to disrupt the CaM mediated activation of eEF2K, presenting both the first *de novo* inhibitor of an alpha kinase and the first inhibitor of eEF2K that does not directly target the active site of the kinase. Our top engineered *de novo* binder, ‘CAM1’, inhibits kinase activity of eEF2K *in vitro*, comparable to the effects of the leading small molecule inhibitor, A-484954. Our *de novo* binder is remarkably thermostable and binds to the CBD of eEF2K with strong affinity. Preliminary studies with model cancer cell lines suggest the transfection of CAM1 reduces cell proliferation and inhibits phosphorylation of eEF2, paving the way for more advanced *in vivo* and translational studies.

## RESULTS

### Computational design of an eEF2K *de novo* binder generates highly accurate hits

eEF2K is a calmodulin activated alpha kinase. Its phosphorylation of eEF2 T56 residues results in the arrest of ribosomal peptide translation. We sought a variety of different strategies when designing a *de novo* binder that targets eEF2K including binding to the eEF2 binding domain, alpha kinase domain, or another site. However, because of the unique factor that eEF2K is CaM activated, and CaM activation has been demonstrated to increase eEF2K activity 100-fold, our binder design strategy was to bind the CaM binding domain of eEF2K, targeting residues 78-105 (PDB: 7SHQ)^1^ (Figure 1b-c). The CBD of eEF2K contains a 1-5-8-14 hydrophobic motif and has an ordered secondary structure (alpha-helix) which meant it was an ideal candidate for binder design with RFDiffusion^1^. Furthermore, this site contained Pro98. When hydroxylated, Pro98 may impair the CaM-eEF2K interaction, thus we realized the target site for RFDiffusion binder design was critical to eEF2K activity and had a reasonable chance of enzymatic inhibition when targeted^15^.

Previous structural determination of the eEF2K-CaM complex has identified that the predominant driver of the CBD-CaM interaction was hydrophobic in nature^1^. Moreover, eEF2K W85 interacted with a deep hydrophobic pocket on CaM^1^. It was found that the CaM C-lobe was ’intimately associated with the kinase’, providing a structural basis for the CaM driven activation of eEF2K^1^. We used RFDiffusion to generate a *de novo* backbone against the CBD of eEF2K. This was done with and without hotspots targeting the 1-5-8-14 hydrophobic motif on the CBD, as we sought to generate a protein-binder interaction partially driven by the hydrophobic effect. 8 binder designs were generated from each RFDiffusion run and for each of these binders, 8 sequences were predicted with ProteinMPNN, resulting in 64 sequences for sequence structure prediction using AlphaFold. The total time to generate a lead *de novo* binder candidate was under one day (Figure 1c).

We sorted CBD binders by AlphaFold PLDDT (per residue confidence score, higher the better) and PAE (predicted aligned error, to assess domain packing and topology, lower the better) output scores. Of 64 binder backbone-sequence pairs, 11 had a PAE <10 and PLDDT >0.9 (Figure 1d). We selected two binders with the lowest PAE scores (<6), which had PLDDT scores >0.92, that are named CAM1 and CAM2. These binders target residues 78-105 of the CBD but RFDiffusion was not configured to target specific hotspots.

Resulting binders CAM1/2 are 105 amino acids in length, have the same predicted backbone, but are fitted with different sequences (Figure 1e). A BLASTP search of these binder sequences returned no related sequences, unsurprising to us as these are *de nov*o sequences. The CAM1/2 structure is a four-membered helical bundle that interacts closely with the helical CBD, together, taking on a helical bundle (Figure 1f). Alignment of CAM1 and CAM2 revealed several regions of sequence difference, throughout the protein. Noticeably, ProteinMPNN did not produce sequences that differed by more than 4 consecutive amino acids at any given sequence position. We next analyzed the protein-protein contacts, applying an 8 Å cutoff between backbone alpha carbons, of the CBD of eEF2K and CAM1/2 (Figure S1b-c). CAM1/2 both come in close contact with several aromatic hydrophobic residues on the CBD including Phe81, Trp85, Trp99, and Phe102, which may be a strong predictor that a hydrophobic interaction between binder and target would occur. Interestingly, for CAM1/2 several contacts are made with Pro98 of the CBD, where the hydroxylation of this residue has been demonstrated to impact EEF2K activity^15^. It is worthwhile to note that while nearly all residues of the CBD are predicted to contact the CAM1/2 binders, only a few helices of the four helical bundle CAM1/2 contact the CBD.

### The *de novo* eEF2K binder, CAM1, inhibits kinase activity *in vitro*

Codon optimized CAM1 and CAM2 suitable for recombinant expression in *E. coli* was commercially synthesized with an N-terminal 6xHis tag and incorporated into the pET28a-(+) vector. Subsequent recombinant purification of CAM1 and CAM2 by nickel affinity chromatography followed by size-exclusion chromatography resulted in highly purified protein (Figure 1g and S2d). Of note, CAM1 and CAM2 produced remarkable yields, always over 5 mg recombinant protein/per culture liter amongst several independent purifications.

To assess the effects of our *de novo* binders on eEF2K activity we used recombinantly purified eEF2K, CaM, and eEF2 peptide substrate (N-RKKYKFNEDTERRRFL-C), using an ADP detection universal kinase assay (Figure S2a-c,e). Titration of the eEF2 peptide substrate resulted in an increase in detected fluorescence, demonstrating the substrate dependent activity of our recombinant eEF2K (Figure 2c). Loss of activity was also demonstrated by titration of EDTA in the reaction mixture, which is expected to chelate calcium ions, resulting in the inactivation of Ca-CaM mediated eEF2K activity (Figure 2b). Triplicate measurements of reactions without either ATP, peptide substrate, or CaM, demonstrated all are required for the kinase reaction and maximum detectable activity of eEF2K (Figure 2a).

**Figure 2.**
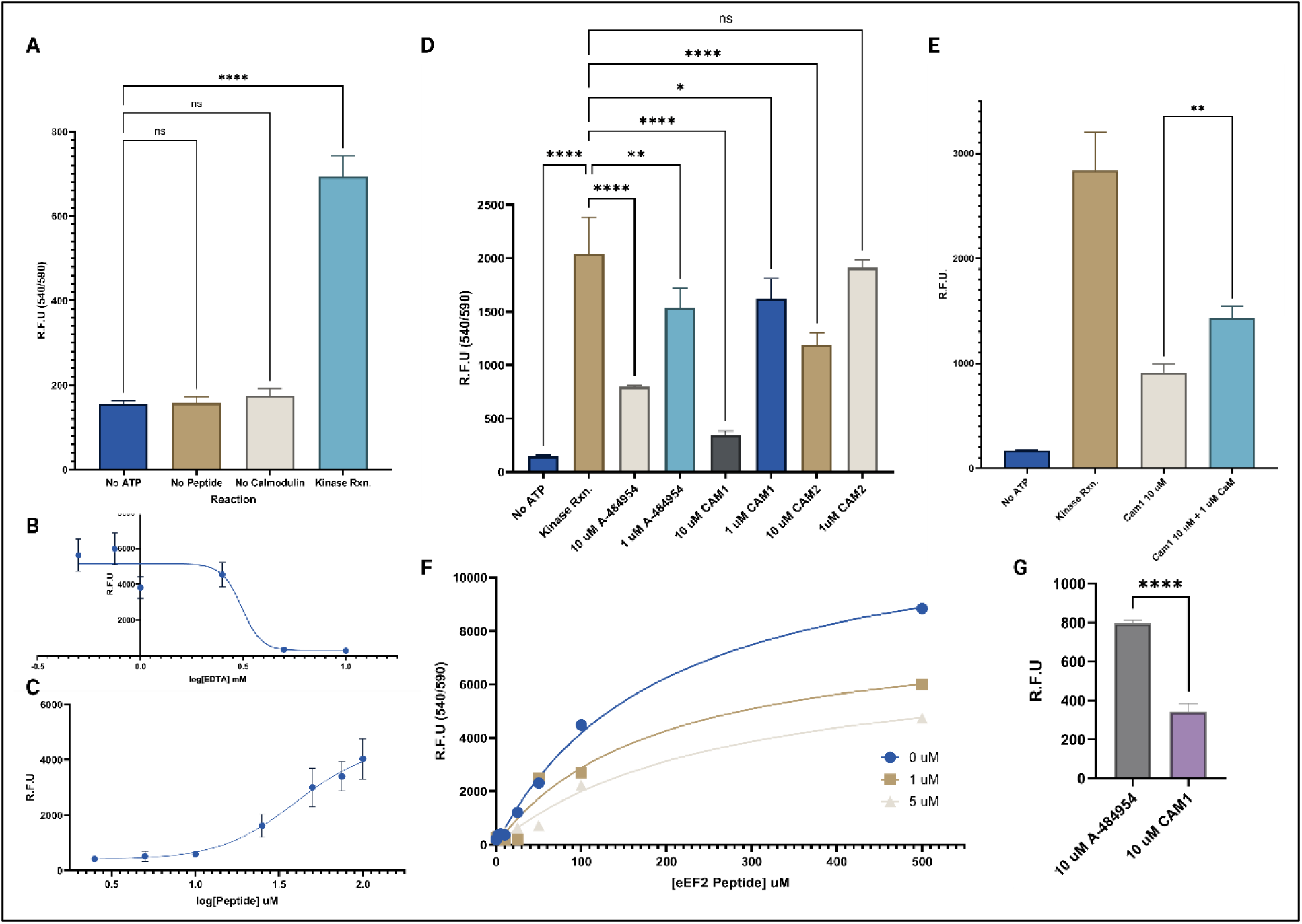
CAM1 inhibits kinase activity of eEF2K. eEF2K kinase activity is assessed by *in vitro* enzymatic kinase assay with eEF2 peptide substrate. A) Compared to no ATP control, eEF2K kinase activity is significantly increased with ATP, peptide substrate, and calmodulin in reaction system (p < 0.0001). B) eEF2K kinase activity is sensitive to addition of EDTA (R^2^ = 0.8718) and is C) stimulated by presence of peptide substrate (R^2^ = 0.9272). D) No ATP, 10 uM of CAM1, CAM2, and A-484954 all significantly inhibit kinase activity (p<0.0001, for all) alongside only 1 uM of CAM1, A-484954, but not CAM2 (p<0.0373, p<0.0100, and p<0.9457, respectfully). E) Compared to 10 uM of CAM1 alone, addition of 1 uM CaM significantly rescues kinase activity but not to baseline (p = 0.0030). F) CAM1 0, 1, and 5 uM series over eEF2 peptide substrate titration reveals a likely mixed inhibitor model (Vmax/Km/R^2^ = 12360/195.3/0.9951, 8263/187.0/0.9556, and 7280/265.1/0.9737 for 0, 1, and 5 uM, respectively). G) Compared to the gold standard small molecule eEF2K inhibitor, the protein binder CAM1 is significantly more potent at equimolar (10 uM) screening concentrations (p<0.0001).

We replicated our initial kinase test conditions with the addition of purified *de novo* binder CAM1/2 or A-484954, an eEF2K specific small molecule inhibitor. eEF2K kinase activity *in vitro* was significantly inhibited by 10 uM A-484954, 10 uM CAM1, 10 uM CAM2 (p = <0.0001, <0.0001, <0.0001, respectively) and at 1 uM of inhibitor was significantly inhibited by A-4984954, CAM1, but not CAM2 (p = 0.01, 0.0373, and 0.9457, respectively) (Figure 2d). As such we narrow the focus for future experiments using CAM1, which reduced kinase activity greater than the leading small molecule drug, A-484954 at equimolar screening concentration (Figure 2g). Despite sharing a similar predicted backbone, differences in inhibition between CAM1/2 may be a result of minor sequence differences produced by ProteinMPNN, however, the extent of these differences on activity and more importantly, which residues impact eEF2K inhibition through CBD binding, remains uncertain.

Since CAM1/2 are designed to bind the CBD of eEF2K and disrupt CaM binding, we sought to test this feature of our binding protein with the addition of excess CaM in the kinase assay. In the presence of 10 uM CAM1, 1 uM of excess CaM was able to rescue some kinase activity for eEF2K (Figure 2e). However, in the CAM1 sample, CaM addition was unable to return eEF2K near the baseline kinase activity. Results suggest the direct targeting of the CBD with the *de novo* binder and competition with CaM for binding of this site which is required for kinase activation.

To highlight that CAM1 is a CBD binder and does not bind at the active site of the eEF2K kinase domain, like the small molecule inhibitor A-484954, we titrated eEF2 peptide substrate under varying CAM1 concentrations (0, 1, and 5 uM) (Figure 2f). With increasing CAM1 concentrations Vmax was reduced, however, Km was relatively unchanged, likely following a mixed model of inhibition. To our knowledge this is the first inhibitor of eEF2K which is not designed to be competitive or directly bind the kinase active site.

### CAM1 is remarkably thermostable

Previous studies have identified the role of three key CBD eEF2K residues His80, His87, His94 (which are included in our binder’s target region), as sensitive to changes in pH^30^. These histidine’s when protonated, are thought to increase the activation of eEF2K by CaM binding^30^. At the structural level it remains unclear how CBD His-protonation may promote the binding of CaM. Since our *de novo* CAM1 is designed by RFDiffusion and ProteinMPNN, this may not consider the effects of histidine protonation on CBD-CAM1 interaction. To our surprise, inhibition was retained across pH range 6-9 (Figure 3a). However, at higher pH (>7.5) we see greater inhibition with 10 uM of CAM1 added to the kinase reaction. These results demonstrate the stability of the CAM1 protein and its inhibitory effects over an extended pH range which may be well suited for future translational work, as the acidic microenvironment of the tumor is below physiological pH^30^.

**Figure 3.**
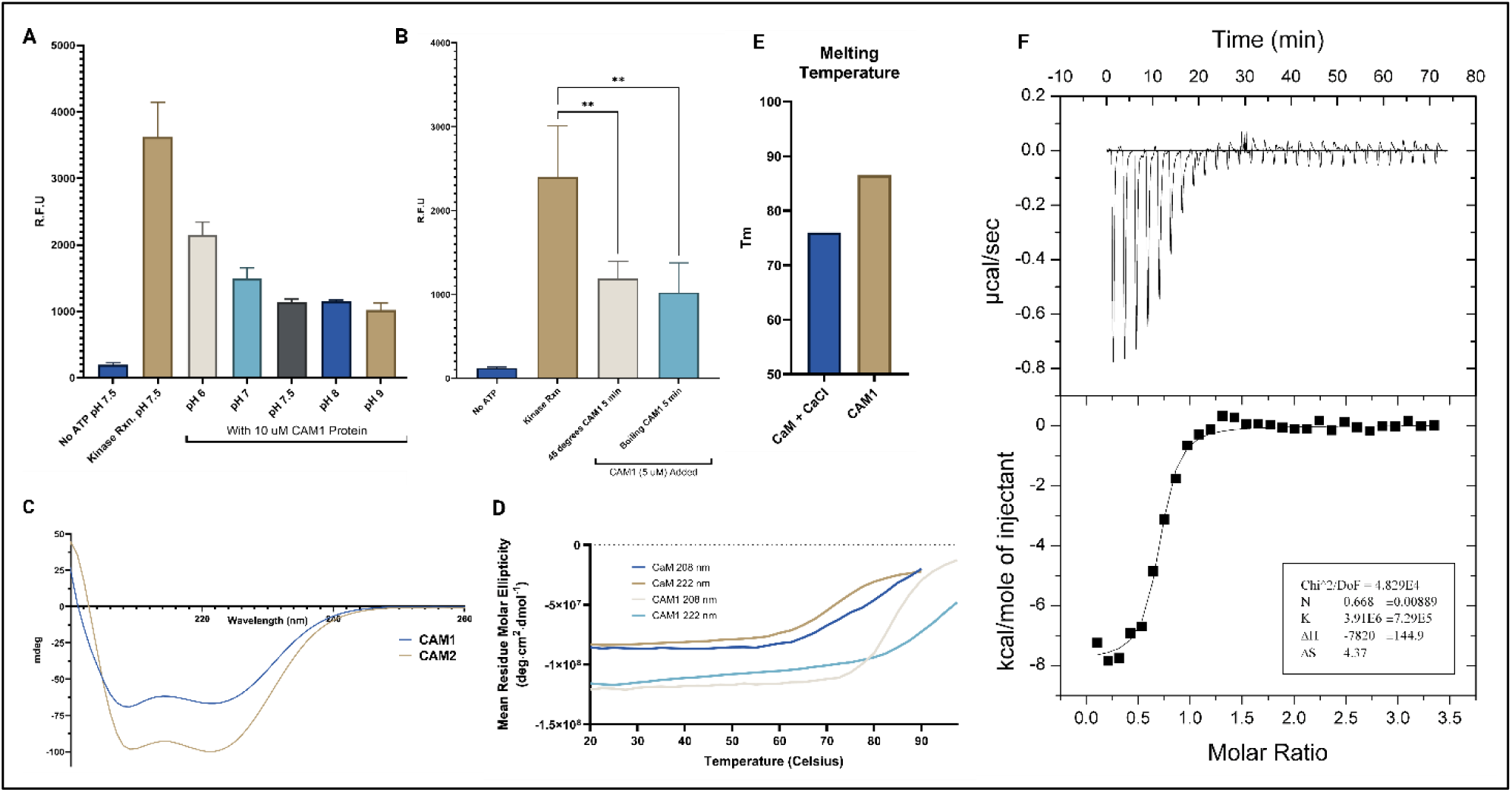
CAM1 is folded, stable, and binds the CBD of eEF2K with high affinity. A) eEF2K kinase activity remains significantly inhibited across a pH gradient (pH 6 to 9) with 10 uM CAM1 protein (one- way ANOVA, p<0.0001). B) 5 uM CAM1 subjected to thermal stress (45°C or boiling for 5 minutes) still inhibits eEF2K kinase activity (p = 0.0321 and 0.0283 for 45°C or boiling, respectively). C) Circular dichroism spectroscopy (CD) spectra of purified CAM1 and CAM2 indicates folded protein with rich alpha helical content. D) CD denaturation study demonstrates high thermostability of CAM1 above the melting temperature of CaM. E) Melting temperature of CAM1 is greater than CaM. F) Representative isothermal titration calorimetry (ITC) binding curve between CBD of eEF2K and CAM1, exothermic reaction which has a binding constant (K) of ∼255 nM.

To assess the thermostability of our CAM1 binder we subjected protein sample to heat stress (45°C or boiling for 5 minutes) and subsequently assessed kinase activity. Much to our surprise significant inhibition of the kinase was still retained with 5 uM of CAM1 treated with either level of heat stress (Figure 3b).

The predicted structure of CAM1/2, contains a high percentage of alpha helical secondary structure content, and a few shorter intrinsically disordered loops. We sought to confirm this by circular dichroism spectroscopy (CD) which assesses the secondary structural content and folding of protein samples. As expected, the CD spectrum of recombinantly purified CAM1 and CAM2 represent a sample with a high percentage of folded alpha helices, as demonstrated with the absolute peaks at approximately 208 and 222 nm, respectively (Figure 3c). Despite small differences in sequence, it is predicted that the tertiary fold and secondary structural elements are near equivalent for both CAM1 and CAM2. CD spectra of CaM had an enriched absolute peak at 208 and 222 nm, alpha helical regions when in the presence of calcium (Figure S3c). Addition of EDTA removed this change in helicity, further confirming the presence of a functionally active CaM (Figure S3c).

We assessed the thermal stability of CAM1 using CD, applying a melting denaturation curve between 20 and 97.5 °C. (Figure 3d, S3b) Alpha helical rich content of the protein sample was also detected by CD using this method. The approximate melting temperature (50% fraction unfolded), Tm, of CAM1 is estimated to be above 85°C, and confirms the thermostable characteristic of this alpha helical rich bundle domain. CaM was also subjected to CD melting denaturation between 20 and 90°C, which had a Tm of over 70°C, particularly lower than that of CAM1 (Figure 3d-e, S3a).

### CAM1 interacts with the eEF2K calmodulin binding domain with nanomolar affinity

Binding of CaM in the presence of calcium to the CBD of eEF2K was evaluated using isothermal titration calorimetry. As expected, binding did occur for the native CaM-CBD interaction (Figure S4b). Next, we determined the affinity of the *de novo* CAM1 for the eEF2K CBD with ITC. Titration of CBD peptide into CAM1 sample demonstrated high affinity (binding constant K of 255 nM) with an exothermic reaction and confirmed that enzymatic inhibition of eEF2K mediated by CAM1 is likely due to the direct binding on the CBD target (Figure 3f).

CaM mediated activation of eEF2K is calcium dependent. Using ITC, we titrated free calcium into CAM1 protein sample and did not detect the presence of binding (Figure S4a). This confirms that our *de novo* binder is designed to only bind the CBD of eEF2K and does not compete with CaM for free calcium.

### The *de novo* CAM1 restricts proliferation and inhibits eEF2K activity in cancer cells

To assess the *in vivo* effects of CAM1 mediated eEF2K inhibition we compared the effects of CAM1 transfection and treatment with the small molecule drug, A-484954. CAM1 DNA optimized for human expression was cloned into the pCDNA3.1 vector which we refer to as pCDNA3.1-CAM1.

We used the WST-8 assay to monitor HeLa cell proliferation after treatment with 100 uM A-484954 or pCDNA3.1-CAM1 transfection. Compared to untreated HeLa cells that significantly proliferated over 48 hrs, A-484954 treatment and CAM1 transfected cells had their proliferation restricted and no noticeable difference was observed (Figure 4a-b). We next sought to assess the anti-proliferative properties of CAM1 transfection by lowering the mass of transfected DNA to 10 ng or 50 ng per 96-well plate well, one tenth or one half of the reagent manufacturers recommended DNA mass for transfection, respectively. To our surprise, after 48 hrs, CAM1 cells transfected with one tenth the normal working DNA mass had proliferation restricted and no noticeable proliferative difference was observed (Figure 4c-d).

**Figure 4.**
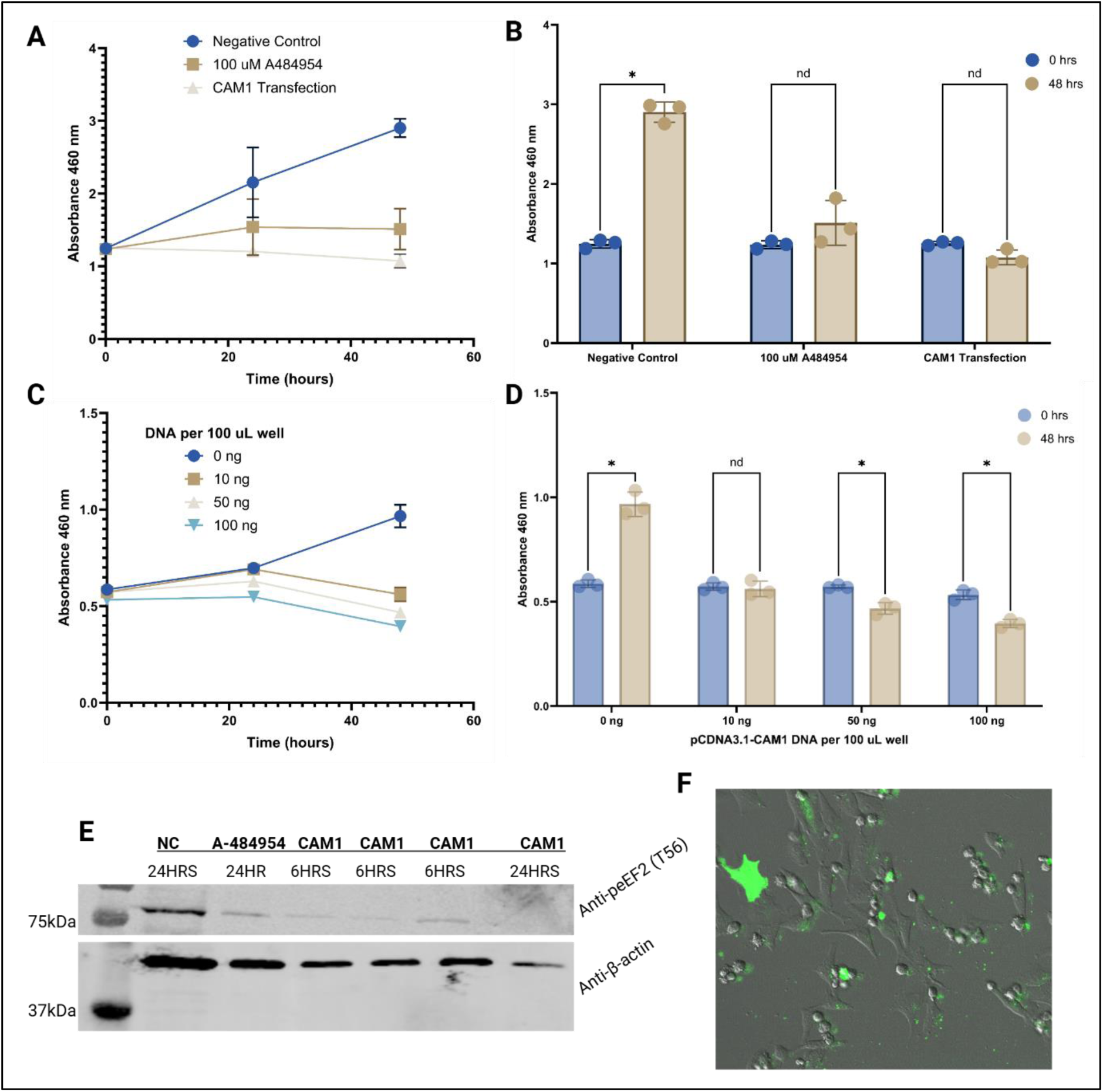
CAM1 inhibits HeLa cell proliferation and eEF2 phosphorylation. A-B) Cell viability WST- 8 assay over 48 hrs with HeLa cells demonstrates cell proliferation is restricted with 100 uM A-484954 and pCDNA3.1-CAM1 transfection but significantly proliferates when untreated (p = 0.000031). C-D) Cell proliferation is restricted by pCDNA3.1-CAM1 transfection between 10 ng and 100 ng per 100 uL volume well of cell-media mixture. E) HeLa cells treated with 100 uM A-484954 for 24 hrs or transfected with CAM1 for 6 plus hrs have a reduction in eEF2K activity as demonstrated with anti-p-eEF2 Thr56 western blotting. F) HeLa cells 6 hrs after transfection of pCDNA3.1-CAM1-GFP.

We next monitored the effects of CAM1 mediated inhibition on eEF2K activity in HeLa cells by probing substrate phosphorylation by western blotting with an anti-peEF2 Thr56 antibody. Similar to cells treated with 100 uM of A-484954 for 24 hrs, CAM1 transfection for either 6 or 24 hrs resulted in a reduction in phosphorylated eEF2 levels in HeLa cells, indicating our inhibitor likely works in a basic *in vivo* cell model to reduce kinase activity (Figure 4e). It is worth noting that at 24 hrs post-transfection, significant cell death was observed and may account for a weaker anti-β-actin signal, nonetheless peEF2 Thr56 signal was still dramatically reduced (Figure 4e). Transfection was also confirmed by conjugation of a GFP tag to the C-terminal of CAM1, generating a pCDNA3.1-CAM1-GFP vector, and expression was confirmed by fluorescent microscopy 6 hrs post-transfection (Figure 4f).

## DISCUSSION

For several decades, small molecule structure-based drug design has been a leading effort for the selective inhibition of eEF2K, amongst other atypical kinases^5^. Despite early successes such as the development of small molecule A-484954, chemical modifications to increase potency *in vivo* require significant expenditures, and are labor intensive^5,22,31^. We have demonstrated that *de novo* protein design for the site directed inhibition of eEF2K may be a viable alternative strategy to targeting this unique, CaM dependent kinase. To our knowledge this is the first *de novo* alpha/atypical kinase protein inhibitor and the first non-competitive eEF2K inhibitor. Several examples of *de novo* protein kinase inhibitors/modulators are present in literature, including *de novo* protein switches designed to induce phosphorylation upon binding a kinase such as Src kinase^32,33^. Due to recent advancements in computational methods such as RFDiffusion, that improve target driven binder design we expect more *de novo* protein kinase inhibitors to appear^28^.

Selective kinase inhibition, amongst atypical kinase families and conventional kinase families remains an ongoing challenge. With respect to conventional protein kinases, targeted inhibition of one kinase amongst hundreds of similarly folded active sites is a prevalent challenge with small molecules, due to high structural conservation in these regions^6,7^. As was the approach in our targeted design of an eEF2K binder, conventional protein kinases with regions of structural dissimilarity such as allosteric domains or substrate binding motifs could serve as good sites for *de novo* binder design. The time to lead candidate with *de novo* binder design as demonstrated herein is markedly lower than time to lead candidate with small molecule inhibitors. We developed our lead binders that were computationally generated and scored in about one week.

We were initially surprised by the high success rate of our binder design strategy and resulting experimental characterization. However, several considerations are worth sharing that we believe improved hit rate and are also similarly found in literature^28^. We selected the CBD of eEF2K as a target not only to disrupt CaM mediated activation of the kinase, but because of its structural features. These include an ordered alpha helical motif and hydrophobic domain. By targeting an ordered secondary structure for binder design, we eliminated any structural promiscuity seen in intrinsically disordered loops and built the binder on a more rigid target. Furthermore, we sought a hydrophobic motif (1-5-8-14 CBD hydrophobic motif) to stimulate the hydrophobic effect of protein-protein interactions, should CAM1/2 find the CBD as expected. Resulting binder designs of CAM1/2 were an alpha helical bundle. It is worth noting that together with the CBD, the binder-target pair is predicted to take on an even larger alpha helical bundle. We presume that this is also a driver of the high success rate through our design. Moreover, we developed a *de novo* binder that is just over 100 amino acids. This is critical as proteins generated with *de novo* design programs and AlphaFold may experience drop off in predicted structural accuracy over several hundred amino acids in size^28,34^.

Inhibition of eEF2K *in vitro* was comparable to the small molecule inhibitor A-484954 when screened at equimolar concentrations in our kinase assay with peptide substrate. While the binding constant was ∼255 nM between the CAM1 and CBD as determined by ITC, it is worth noting that while demonstrating reasonable affinity, kinase inhibition and binding can be improved in the future with either site directed modification of the lead CAM1 binder or partial diffusion technique^28^. With partial diffusion a similar backbone structure can be generated alike CAM1/2, but after applying a new sequence, may exhibit greater potency and affinity for the CBD. Models using RFDiffusion in literature for binder design, demonstrated binding affinity in the very low nanomolar range, thus we predict that with greater construct design throughput this is readily achievable for the case of an eEF2K binder^28^.

The stability of our CAM1 binder was also comparable to other *de novo* designed small proteins. Considering that CAM1 is predicted to be an alpha helical bundle, its remarkable thermostability and inhibition after heat treatment are both unsurprising to us. Considering the inhibition of the kinase is retained with CAM1 across a pH gradient, and below neutral pH, this may be suitable for an acidic microenvironment like tumors. We did not consider when designing a CBD binder, the effects of pH on ionizable residues in the CBD such as histidine’s, or the hydroxylated Pro98 residue. Such design implementations would be particularly interesting in subsequent work to see if a *de novo* binder could be designed to demonstrate inhibition only at non-physiological pH.

Final characterization of CAM1 for this work involved assessment and the positive confirmation of eEF2K inhibition in a cancer cell line (HeLa cells). Reduction of p-eEF2 Thr56 phosphorylation is consistent with other studies using small molecule eEF2K inhibitors such as A-484954^22^. We also noticed an arrest in HeLa cell proliferation with CAM1 transfection, similar to cells treated with 100 uM of A- 484954, and furthermore this arrest in cell proliferation was present at even 1/10^th^ the recommended DNA transfection levels. CAM1 expression and potency at just 6 hrs post-transfection was particularly interesting to observe. While CAM1 design involved specific structure driven binder design, off target effects at this time remain unknown. Since the CBD of eEF2K is dissimilar to other motifs found in the human proteome specificity of CAM1 to its target may be greater than other conserved kinase motifs.

Future experiments seek to understand how CAM1 may affect proliferation and eEF2 phosphorylation in other cell lines, in both disease cell lines such as cancer, and healthy cells. In future work we seek to test CAM1 more extensively *in vivo*, targeting the *de novo* binder to cancer cells and thus removing the uncertainties of ill effects with CAM1 mediated eEF2K inhibition in healthy cells. Such experiments rely heavily on delivery systems for protein-based therapeutics which are more challenging to deliver to cytosolic targets than small molecule inhibitors, primarily due to molecular weight and size. Solutions to this problem may be mRNA-based delivery and inclusion of a cancer specific gene promoter, such as hTERT, to restrict the expression of CAM1 to only cancer cells^35^.

## METHODS

### *De Novo* backbone design and sequence prediction

To generate binders against the CBD of eEF2K, we used a CaM-eEF2K experimentally determined structure (PDB: 7SHQ) for target input. RFDiffusion was used to develop a *de novo* binders’ backbone with a favorable fit to the CBD site and was configured to design a binder between 75-150 amino acids in size, against residues 78-105 of the CBD. We configured RFDiffusion to perform 50 iteration steps for each of the 8 binder designs.

For each of the 8 backbone designs, we used ProteinMPNN to generate 8 sequences (total designs = 64) to place appropriate side chains that would support this protein fold and interaction with the CBD of eEF2K. Predicted sequences were passed to AlphaFold2 which predicted the structure of the *de novo* binder by sequence only.

### Scoring of *de novo* protein designs

*De novo* binders were assessed by AlphaFold2 scores PAE (predicted aligned error) and PLDDT (predicted local distance difference test). We considered binders with a PAE <10 and PLDDT >0.9 for further analysis as these binders have the highest likelihood of a sequence that folds into the predicted backbone structure. CAM1/2 binders are the two designs with the lowest PAE scores.

### Bioinformatic search of CAM1/2 sequence and interaction with CBD

Protein sequences for CAM1/2 were inputted into BLASTP and compared against all non-redundant protein sequences in the standard databases^36^. No organisms were filtered or removed. Algorithm parameters were set to default with a BLOSUM62 scoring matrix used.

Interactions between CAM1/2 and the CBD were calculated, briefly, as follows. Using the Biopython PDBParser module, distances between the alpha carbon backbone atoms between the two protein chains were calculated. Contacts greater than 8 Å apart were rejected, and those within the threshold were plotted.

### CAM1/2 recombinant protein purification

CAM1/2 sequences appended to an N-terminal 6xHistidine tag were synthesized, commercially, and subcloned into pET28a-(+) vector. Vector was transformed into BL21-RIPL *E. coli* cells with LB-Agar plates containing Kanamycin for selection. CAM1/2 positive colonies were inoculated into 5 mL starter LB cultures at 37°C O/N, then added to 500 mL of LB media, which was grown at 37°C, until O.D 600nm of 0.6, and finally expressed at 20°C O/N.

Cells were pelleted at 4,000G at 4°C for 30 minutes. Pellets were resuspended in approximately 25 mL of lysis buffer (20 mM sodium phosphate, 300 mM NaCl, pH 7.4), sonicated, and clarified with high-speed centrifugation (18,000G at 4°C for 30 minutes). Nickel resin was mixed with supernatant, rocking gently at 4°C for 30 minutes. Low speed centrifugation (1,000G at 4°C for 3-5 minutes) was used to collect CAM1/2 bound resin, which was resuspended in 10 mL lysis buffer, as flow through for gravity column purification. Protein on the resin was eluted with lysis buffer mixed with an imidazole gradient (50-, 75-, 100-, and 200-mM imidazole), at 15 mL total volume for each elution fraction.

Elution fractions were analyzed on SDS-PAGE, and fractions containing CAM1/2 protein were pooled and concentrated by centrifugation. Protein was further purified by size-exclusion chromatography on FPLC system, with a column pre-equilibrated with lysis buffer. Fractions containing protein were analyzed by SDS-PAGE, pooled, and concentrated by centrifugation to approximately 10 mg/mL prior to being flash frozen.

### eEF2K recombinant protein purification

Human eEF2K cDNA clone in pGEM-T cloning vector was purchased commercially (Sino Biological Cat: HG11587-G) and resuspended in 100 uL of sterile water. eEF2K cDNA was amplified and subcloned into a new expression vector with N-terminal 6xHis tag and maltose binding protein (MBP) to promote soluble bacterial expression and purification. The plasmid was transformed into BL21-RIPL *E. coli* cells with LB-Agar plates containing Ampicillin for selection. His-MBP-TEV-eEF2K positive colonies were inoculated into 5 mL starter LB cultures at 37°C O/N, then added to 500 mL of LB media, which was grown at 37°C, until O.D 600nm of 0.6, and finally grown at 20°C O/N. Unlike the purification of CAM1/2, several litres of culture were required for higher purification yields of recombinant eEF2K.

Recombinant eEF2K was purified by nickel affinity chromatography as described with the recombinant purification of CAM1/2. After SDS-PAGE analysis of fractions washed with imidazole pooled fractions containing purified eEF2K were pooled and TEV protease was added (1:100 TEV protease:eEF2K), rocking at 4°C O/N. Protein was further purified by size-exclusion chromatography on FPLC system, with a column pre-equilibrated with lysis buffer. Fractions containing eEF2K protein separated from MBP were analyzed by SDS-PAGE, pooled, and concentrated by centrifugation to approximately 6 mg/mL prior to being flash frozen.

### Calmodulin recombinant protein purification

Gene encoding CaM protein appended to an N-terminal 6xHisidine tag and TEV protease cut site, inserted into the pDest-527 *E. coli* expression vector was purchased from Addgene (Plasmid #159693). Vector was transformed into BL21-RIPL *E. coli* cells with LB-Agar plates containing Ampicillin for selection. CaM was purified using the same methods described for CAM1/2, however, prior to loading onto FPLC, was mixed with TEV protease (1:100 w/w TEV protease:CaM), for several hours, rocking at 4°C. CaM used in enzymatic assays was supplemented with free calcium to enable the calcium-CaM mediated activation of eEF2K.

### eEF2K Kinase Activity Assays

eEF2 phosphorylation site peptide substrate was commercially synthesized (N- RKKYKFNEDTERRRFL-C) for use with an enzymatic kinase activity assay^37^. Recombinantly purified CAM1/2, CaM, and eEF2K were all used for kinase activity assays. Similarly described in literature, we used 6 ng/uL (final concentration) of eEF2K, equimolar CaM, and 1 uM of peptide substrate, and 0.1 mM ATP for all kinase assays^38^. 1 mM of CaCl was added for calcium-CaM mediated activation of eEF2K. A fluorometric Abcam Universal Kinase Assay Kit (product code ab138879) was used and followed according to the manufacturers protocol, which measures the formation of ADP. Kinase reactions were carried out at 37^0^C in buffer containing 40 mM Tris (pH 7.5), 20 mM MgCl2 and 0.1 mg/mL BSA.

For kinase assays in which CAM1 was subjected to heat stress, CAM1 was returned to room temperature prior to addition in the assay. For kinase assays in which inhibition at a given pH was assessed, kinase reaction buffer was pH adjusted.

### Circular dichroism spectroscopy and melting temperature assessment

Circular dichroism spectroscopy (CD) studies were conducted using an Applied Photophysics Ltd. Chirascan V100 spectrophotometer device. Purified protein samples were diluted to approximately 0.5 mg/mL and spectra was collected between 190 and 260 nm. A disc cuvette was used to minimize path length (Hellma Usa Inc., round cylindrical cuvette, 1mm pathlength). For studies with CaM, 1 mM of CaCl or 1 mM of EDTA was added as described.

Melting temperature Tm measurements were collected using CD. Tm is the temperature at which the fraction folded is equivalent to the fraction unfolded and calculated using the equation Fu = (Fo-F)/(Uf- F), where F is the ellipticity value at low temperature, Fo is the value at low temperature, and Uf is the value at the highest (unfolded) temperature). Prior to thermal denaturation protein aliquots were removed from cold storage, quick thawed at room temperature, and stored on ice. The cuvette was placed into the CD spectrophotometer and a water bath/circulator heated the sample.

### Isothermal titration calorimetry

For isothermal titration calorimetry (ITC) studies, a VP-ITC (Malvern Panalytical) was used. Protein or peptide samples were prepared in the same buffer (20 mM sodium phosphate, 300 mM NaCl, pH 7.4). Interactions were analyzed using Origin (MicroCal, LLC) software. For all ITC experiments, temperature was set to 30^0^C, the differential power was set to 25 (uCal/Sec), stirring speed was set to ∼300 rpm, and 10 uL of sample in syringe was injected into the cell every 150 seconds for 30 total injections.

ITC was configured such that the CBD peptide (300 uM) was injected into the cell containing either CAM1 (20 uM) or CaM (30 uM). Separately, 300 uM of CaCl2 was injected into the cell containing either 30 uM of CAM1 or CaM to study the effects of calcium binding.

### Cell proliferation assays

HeLa cells between passage number 3 to 20 were used for cellular studies. HeLa cells were evenly seeded onto a 96-well tissue culture plate and after reaching over 50% confluency, pCDNA3.1-CAM1 was transfected using Lipofectamine3000 kit, according to manufacturers protocol (ThermoFisher) or cells were treated with A-484954 drug (final concentration of 100 uM). WST-8 assay was used according to the manufacturers protocol (Abcam ab65475) and absorbance was recorded between 0 and 48 hrs following transfection. For studies assessing the potency of pCDNA3.1-CAM1 transfection, Lipofectamine3000 reagents were still used at equivalent concentrations, and only the amount of pCDNA3.1-CAM1 was altered.

### Western blot studies

HeLa cells between passage number 3 to 20 were used for cellular studies. HeLa cells were evenly seeded onto a 6-well tissue culture plate and after reaching over 50% confluency, pCDNA3.1-CAM1 was transfected using Lipofectamine3000 kit, according to manufacturers protocol (ThermoFisher) or cells were treated with A-484954 drug (final concentration of 100 uM). After 6 or 24 hrs, cells were lysed as described in Liao et al. in buffer containing 20 mM HEPES, pH 7.5, 50 mM NaCl, 25 mM KCl, 10 mM DTT, 3 mM benzamidine, 1% SDS, 1 mM sodium orthovanadate, 20 mM sodium pyrophosphate, and 1 tablet of cysteine protease inhibitors/10 mL^39^. Antibody against phosphorylated T56 eEF2 (Cell Signaling Technology #2331) was used to assess the inhibition of eEF2K and antibody against beta-actin was used as a loading control (Cell Signaling Technology #3700).

### CAM1-GFP microscopy

HeLa cells between passage number 3 to 20 were used for microscopy studies. HeLa cells were evenly seeded onto a 35 mm collagen coated disk with glass bottom (Mattek Part No P35GCOL-1.5-14-C) and after reaching over 50% confluency, pCDNA3.1-CAM1-GFP was transfected using Lipofectamine3000 kit, according to manufacturers protocol (ThermoFisher). Transfection and cell morphology was assessed at varying time points using live-cell fluorescent imaging (Mica microscope, Leica Microsystems). Brightfield/IMC and GFP fluorescent channels were overlayed post acquisition for visualization purposes.

## ACKNOWLEDGEMENTS

We thank Dr. Laura van Staalduinen (Queen’s University, Canada) for her thought-provoking conversations and expert advice on studying alpha kinases. We thank Matty Graham (Queen’s University, Canada) for his assistance will mammalian cell culture. Cartoon figures were created using BioRender.

## AUTHOR CONTRIBUTIONS

Conceptualization, Z.J., K.K.; Methodology, K.K.; Investigation, K.K, and E.B.; Writing – Original Draft, K.K.; Writing – Review and Editing, K.K., E.B., and Z.J.; Funding Acquisition, Z.J.; Supervision; Z.J.

## REFERENCES

1. Piserchio, A. et al. Structural basis for the calmodulin-mediated activation of eukaryotic elongation factor 2 kinase. Science Advances 8, eabo2039 (2022).

2. Ryazanov, A. G. et al. Identification of a new class of protein kinases represented by eukaryotic elongation factor-2 kinase. Proc. Natl. Acad. Sci. U.S.A. 94, 4884–4889 (1997).

3. Kenney, J. W., Moore, C. E., Wang, X. & Proud, C. G. Eukaryotic elongation factor 2 kinase, an unusual enzyme with multiple roles. Adv Biol Regul 55, 15–27 (2014).

4. Kaul, G., Pattan, G. & Rafeequi, T. Eukaryotic elongation factor-2 (eEF2): its regulation and peptide chain elongation. Cell Biochemistry and Function 29, 227–234 (2011).

5. Klupt, K. A. & Jia, Z. eEF2K Inhibitor Design: The Progression of Exemplary Structure-Based Drug Design. Molecules 28, 1095 (2023).

6. Drennan, D. & Ryazanov, A. G. Alpha-kinases: analysis of the family and comparison with conventional protein kinases. Prog. Biophys. Mol. Biol. 85, 1–32 (2004).

7. Middelbeek, J., Clark, K., Venselaar, H., Huynen, M. A. & van Leeuwen, F. N. The alpha-kinase family: an exceptional branch on the protein kinase tree. Cell. Mol. Life Sci. 67, 875–890 (2010).

8. Ryazanov, A. G., Pavur, K. S. & Dorovkov, M. V. Alpha-kinases: a new class of protein kinases with a novel catalytic domain. Current Biology 9, R43–R45 (1999).

9. Pigott, C. R. et al. Insights into the regulation of eukaryotic elongation factor 2 kinase and the interplay between its domains. Biochem J 442, 105–118 (2012).

10. Pavur, K. S., Petrov, A. N. & Ryazanov, A. G. Mapping the functional domains of elongation factor-2 kinase. Biochemistry 39, 12216–12224 (2000).

11. Piserchio, A. et al. Structural dynamics of the complex of calmodulin with a minimal functional construct of eukaryotic elongation factor 2 kinase and the role of Thr348 autophosphorylation. Protein Sci 30, 1221–1234 (2021).

12. Piserchio, A. et al. Solution Structure of the Carboxy-Terminal Tandem Repeat Domain of Eukaryotic Elongation Factor 2 Kinase and Its Role in Substrate Recognition. J. Mol. Biol. 431, 2700–2717 (2019).

13. Tavares, C. D. J. et al. The Molecular Mechanism of Eukaryotic Elongation Factor 2 Kinase Activation. J Biol Chem 289, 23901–23916 (2014).

14. Liu, R. & Proud, C. G. Eukaryotic elongation factor 2 kinase as a drug target in cancer, and in cardiovascular and neurodegenerative diseases. Acta Pharmacol Sin 37, 285–294 (2016).

15. Moore, C. E. J. et al. Elongation Factor 2 Kinase Is Regulated by Proline Hydroxylation and Protects Cells during Hypoxia. Mol Cell Biol 35, 1788–1804 (2015).

16. Warburg, O. On the origin of cancer cells. Science 123, 309–314 (1956).

17. Cheng, Y. et al. eEF-2 kinase is a critical regulator of Warburg effect through controlling PP2A-A synthesis. Oncogene 35, 6293–6308 (2016).

18. Bircan, H. A. et al. Elongation factor-2 kinase (eEF-2K) expression is associated with poor patient survival and promotes proliferation, invasion and tumor growth of lung cancer. Lung Cancer 124, 31– 39 (2018).

19. Wang, R.-X., Xu, X.-E., Huang, L., Chen, S. & Shao, Z.-M. eEF2 kinase mediated autophagy as a potential therapeutic target for paclitaxel-resistant triple-negative breast cancer. Annals of Translational Medicine 7, 783 (2019).

20. Gschwendt, M., Kittstein, W. & Marks, F. Elongation factor-2 kinase: effective inhibition by the novel protein kinase inhibitor rottlerin and relative insensitivity towards staurosporine. FEBS Letters 338, 85–88 (1994).

21. Arora, S. et al. Identification and characterization of an inhibitor of eukaryotic elongation factor 2 kinase against human cancer cell lines. Cancer Res 63, 6894–6899 (2003).

22. Chen, Z. et al. 1-Benzyl-3-cetyl-2-methylimidazolium iodide (NH125) Induces Phosphorylation of Eukaryotic Elongation Factor-2 (eEF2). J Biol Chem 286, 43951–43958 (2011).

23. Liu, Y. et al. Designing an eEF2K-Targeting PROTAC small molecule that induces apoptosis in MDA-MB-231 cells. Eur J Med Chem 204, 112505 (2020).

24. Anfinsen, C. B. Principles that govern the folding of protein chains. Science 181, 223–230 (1973).

25. Chu, A. E., Lu, T. & Huang, P.-S. Sparks of function by de novo protein design. Nat Biotechnol 42, 203–215 (2024).

26. Brian Kuhlman et al. Design of a Novel Globular Protein Fold with Atomic-Level Accuracy. Science 302, 1364–1368 (2003).

27. Sumida, K. H. et al. Improving Protein Expression, Stability, and Function with ProteinMPNN. J. Am. Chem. Soc. 146, 2054–2061 (2024).

28. J. P. Watson et al. De novo design of protein structure and function with RFdiffusion. Nature 620, 1089–1100 (2023).

29. Dauparas, J. et al. Robust deep learning–based protein sequence design using ProteinMPNN. Science (2022) doi:10.1126/science.add2187.

30. Xie, J. et al. Molecular Mechanism for the Control of Eukaryotic Elongation Factor 2 Kinase by pH: Role in Cancer Cell Survival. Molecular and Cellular Biology 35, 1805–1824 (2015).

31. Liu, Y. et al. Designing an eEF2K-Targeting PROTAC small molecule that induces apoptosis in MDA-MB-231 cells. Eur J Med Chem 204, 112505 (2020).

32. Ham, J. M., Kim, M., Kim, T., Ryu, S. E. & Park, H. Structure-Based De Novo Design for the Discovery of Miniprotein Inhibitors Targeting Oncogenic Mutant BRAF. International Journal of Molecular Sciences 25, 5535 (2024).

33. Woodall, N. B. et al. De novo design of tyrosine and serine kinase-driven protein switches. Nat Struct Mol Biol 28, 762–770 (2021).

34. Jumper, J. et al. Highly accurate protein structure prediction with AlphaFold. Nature 596, 583–589 (2021).

35. Kyo, S., Takakura, M., Fujiwara, T. & Inoue, M. Understanding and exploiting hTERT promoter regulation for diagnosis and treatment of human cancers. Cancer Sci 99, 1528–1538 (2008).

36. Altschul, S. F., Gish, W., Miller, W., Myers, E. W. & Lipman, D. J. Basic local alignment search tool. J Mol Biol 215, 403–410 (1990).

37. Abramczyk, O. et al. Purification and Characterization of Tagless Recombinant Human Elongation Factor 2 Kinase (eEF-2K) Expressed in Escherichia coli. Protein Expr Purif 79, 237–244 (2011).

38. Xiao, T., Liu, R., Proud, C. G. & Wang, M.-W. A high-throughput screening assay for eukaryotic elongation factor 2 kinase inhibitors. Acta Pharmaceutica Sinica B 6, 557–563 (2016).

39. Liao, Y. et al. Paradoxical Roles of Elongation Factor-2 Kinase in Stem Cell Survival*. Journal of Biological Chemistry 291, 19545–19557 (2016).

