## Supplementary Information for "Development of a *De Novo* Protein Binder that Inhibits the Alpha Kinase eEF2K"

**SUPPLEMENTARY INFORMATION AND FIGURES**

**Table 1.** Predicted Protein-Protein Contact Distances of CAM1 with the CBD of eEF2K

| Res # eEF2K CBD | Amino Acid | Res # CAM1 | Amino Acid | Distance (Angstroms) |
|---|---|---|---|---|
| 81 | PHE | 99 | ILE | 6.216543 |
| 81 | PHE | 102 | LEU | 7.987834 |
| 81 | PHE | 103 | ALA | 6.711651 |
| 84 | ALA | 99 | ILE | 7.361515 |
| 85 | TRP | 8 | MET | 7.623988 |
| 85 | TRP | 96 | MET | 6.557441 |
| 85 | TRP | 99 | ILE | 6.712446 |
| 88 | ALA | 92 | LEU | 6.636349 |
| 88 | ALA | 95 | GLU | 7.40267 |
| 88 | ALA | 96 | MET | 6.437203 |
| 89 | ILE | 12 | LEU | 7.589449 |
| 89 | ILE | 96 | MET | 7.337105 |
| 91 | LYS | 92 | LEU | 7.899425 |
| 92 | ALA | 89 | TRP | 7.279716 |
| 92 | ALA | 92 | LEU | 6.577487 |
| 92 | ALA | 93 | PHE | 7.920609 |
| 93 | LYS | 15 | LEU | 7.462075 |
| 95 | MET | 89 | TRP | 7.636022 |
| 96 | PRO | 85 | LEU | 7.643504 |
| 97 | ASP | 19 | ARG | 6.701142 |
| 98 | PRO | 82 | GLN | 6.755048 |
| 98 | PRO | 85 | LEU | 6.031095 |
| 98 | PRO | 86 | ILE | 5.734955 |
| 98 | PRO | 89 | TRP | 7.451612 |
| 99 | TRP | 19 | ARG | 7.50106 |
| 99 | TRP | 23 | LEU | 7.382597 |
| 99 | TRP | 67 | HIS | 7.806004 |
| 102 | PHE | 22 | SER | 7.518101 |
| 102 | PHE | 23 | LEU | 6.41384 |
| 102 | PHE | 70 | LEU | 7.742859 |
| 102 | PHE | 71 | ALA | 7.969306 |
| 103 | HIS | 22 | SER | 4.810674 |
| 103 | HIS | 23 | LEU | 5.730586 |
| 104 | LEU | 22 | SER | 6.40494 |
| 104 | LEU | 23 | LEU | 6.926288 |
| 104 | LEU | 24 | PRO | 7.453005 |

**Table 2.** Predicted Protein-Protein Contact Distances of CAM2 with the CBD of eEF2K

| Res # eEF2K CBD | Amino Acid | Res # CAM2 | Amino Acid | Distance (Angstroms) |
|---|---|---|---|---|
| 81 | PHE | B99 | VAL | 5.919204 |
| 81 | PHE | B103 | LEU | 7.938284 |
| 84 | ALA | B99 | VAL | 7.369602 |
| 85 | TRP | B95 | GLU | 7.801312 |
| 85 | TRP | B96 | MET | 6.033175 |
| 85 | TRP | B99 | VAL | 6.523797 |
| 88 | ALA | B92 | LEU | 5.685178 |
| 88 | ALA | B93 | TYR | 7.903357 |
| 88 | ALA | B95 | GLU | 6.826689 |
| 88 | ALA | B96 | MET | 6.454918 |
| 89 | ILE | B12 | LEU | 7.74228 |
| 89 | ILE | B92 | LEU | 7.537046 |
| 89 | ILE | B96 | MET | 7.380319 |
| 91 | LYS | B92 | LEU | 7.693471 |
| 92 | ALA | B89 | TRP | 6.9854 |
| 92 | ALA | B92 | LEU | 6.506395 |
| 93 | LYS | B15 | LEU | 7.591633 |
| 96 | PRO | B85 | LEU | 7.287748 |
| 97 | ASP | B19 | ARG | 6.414878 |
| 97 | ASP | B85 | LEU | 7.710161 |
| 98 | PRO | B82 | GLN | 6.112323 |
| 98 | PRO | B85 | LEU | 5.215586 |
| 98 | PRO | B86 | ILE | 5.452711 |
| 98 | PRO | B89 | TRP | 7.304476 |
| 99 | TRP | B19 | ARG | 7.180788 |
| 99 | TRP | B20 | LEU | 7.395762 |
| 99 | TRP | B23 | LEU | 7.635077 |
| 99 | TRP | B67 | ALA | 7.843782 |
| 99 | TRP | B82 | GLN | 7.862003 |
| 102 | PHE | B23 | LEU | 6.582028 |
| 102 | PHE | B70 | LEU | 7.053145 |
| 102 | PHE | B71 | ALA | 7.186074 |
| 102 | PHE | B74 | LEU | 7.896431 |
| 103 | HIS | B22 | SER | 5.07351 |
| 103 | HIS | B23 | LEU | 5.524511 |
| 104 | LEU | B22 | SER | 6.671568 |
| 104 | LEU | B23 | LEU | 6.967528 |
| 104 | LEU | B24 | PRO | 7.671581 |

**Table 3.** ProteinMPNN Sequence Outputs and AlphaFold Scores

| Design | PLDDT | PAE (Å) | RMSD (Å) | Sequence |
|---|---|---|---|---|
| 0 | 0.81 | 15.58 | 2.96 | SFHFKEAWKHAIQKAKHMPDPWAEFHLE/SALLEDIAAG FSRLLEDLLERLRPLLERFPGDEAAWAAFLAQLRALAEEL GLPPEQIAALLALAEARLAEIRARGAAHPGGSAPVLAQIS ELLQAFAEEVKKLLK |
| 1 | 0.61 | 22.42 | 8.79 | SFHFKEAWKHAIQKAKHMPDPWAEFHLE/SELKEDIKEAI KRLLRDLFEMLRPLLIRTPGDEEARKRFLEEVRELLEELG LDEEQIEEILEEVEAILAEIEAEGAENPGGNEPVLKQIEELV REAAEEIKKILE |
| 2 | 0.87 | 8.79 | 2.59 | SFHFKEAWKHAIQKAKHMPDPWAEFHLE/SSLKEQIKEG VKELIEELFDMYRPLLIRFPGDEEAEEAFLKELRELMEEL GLDEEQIDEILALVRERLAEVREEGAEHPGGNAPVLEQIK EVLREFAEEIKKILK |
| 3 | 0.76 | 21.00 | 9.87 | SFHFKEAWKHAIQKAKHMPDPWAEFHLE/MEIEEDIKETI KRVLLDIFDRLRPLLVRFPGDEEARRRLLEEIRKMLKELG LSEEQIEEILELVEARLAEVRAEGAEHPGGSGPVLEQILEI LKEAVEEIKKILK |
| 4 | 0.85 | 12.98 | 2.00 | SFHFKEAWKHAIQKAKHMPDPWAEFHLE/MKIKEDIKET LERVLKDLFEMYRPLLIRFPGDEEAEKELLENIRKMMKEL GLSEEEIEEILKLTEEILEEIREEGKDNPGGNAPVLKQIQEI LEEAVKEIKKILK |
| 5 | 0.54 | 22.56 | 9.43 | SFHFKEAWKHAIQKAKHMPDPWAEFHLE/EEIKKDIAEAI KRLLLDLFEMRRPLLIRFPGDKEAREKLKKHVEELMKEL GLSEELIKEILEELEKILAEIEKEGAANPGGNKPVLEQIKEI LEEAVEEIKKILE |
| 6 | 0.77 | 13.78 | 2.37 | SFHFKEAWKHAIQKAKHMPDPWAEFHLE/EELLEDIAEG VKRVLLDALEMLRPLLIRFPGDEAARAALLAELEAQLKE LGLPEELIAEVLALLEARLAEIAAEGAEHPGGNAPVLAQI AEILKEFVEEVKKILK |
| 7 | 0.73 | 21.97 | 21.73 | SFHFKEAWKHAIQKAKHMPDPWAEFHLE/SSTLEDIKETI SRLLDDLLEMYRPLLERFPGDEAAEKALLEALRAQLKEL GLPEEQIKEILERLERRLEEIRARGKENPGGSGPVLKQIQE VLEEAVEEIKKILE |
| 8 | 0.91 | 6.79 | 3.72 | SFHFKEAWKHAIQKAKHMPDPWAEFHLE/NEELIEEMEK ELEELIEEFLKSLPEDVAKEVKEAVEKTKEELEKNPLTPEF IEKLKKKLEEEFLKIHEKLAKRLGKEVTPEQVELIREWVK LLMEMLVVIKLLKD |
| 9 | 0.91 | 6.84 | 1.47 | SFHFKEAWKHAIQKAKHMPDPWAEFHLE/NEAEIKELEE ELEKLIEEFLASLPPEAAAEVRAIVEATRRELREAPLTPER VAALRDRLLAELLALAERLAARLGRALTPRQRELIAEWV DLLMEMLTVIKLLEL |
| 10 | 0.93 | 5.79 | 1.41 | SFHFKEAWKHAIQKAKHMPDPWAEFHLE/KEEEIEEMEE ELEELIEKRLASLPPEVREKLKAIIEETKKLLEENPLTPEFE |

| | | | | |
|---|---|---|---|---|
| | | | | KELKEKLLKKFLEILKELAKELGKEVTEEQVKLIEEWVELLMTMMVVIKKAKL |
| 11 | 0.91 | 6.76 | 1.84 | SFHFKEAWKHAIQKAKHMPDPWAEFHLE/REEEIEELEKELEELIEERLKSLPPEAAAKLRAIFEKARKKLEENELTPEFVEELKKEVLEEVLALSEELAKKLGKEVTEEQVKLIKEWVELIMEMLVVIKKLKR |
| 12 | 0.94 | 5.25 | 2.66 | SFHFKEAWKHAIQKAKHMPDPWAEFHLE/NEEEIEEMKEELEELLEERLASLPPEVAEEVRRVFEETRERLKKSPLTPERVEELREEMLERLLEIHRRLAAELGREVTPEQVELIRQWVELFIEMLVIIKLAED |
| 13 | 0.93 | 5.52 | 1.81 | SFHFKEAWKHAIQKAKHMPDPWAEFHLE/NEEEIKELKEELEELLEERLKSLPPEVREEILRIYEEARELLKKNPLTPEFIEELKARLLAELLAVAERLAAELGRAMTPEQRALIEEWVELYTEMLVVIKKLEL |
| 14 | 0.94 | 5.88 | 2.24 | SFHFKEAWKHAIQKAKHMPDPWAEFHLE/REEEIAEMEAELDALVAERLASLPPEVAARVRAALERARKRLKENPLTPENVKAIEEELLAELLAISRELAARLGKELTPEQVELIEEWVHQLMTMMTVIELAKR |
| 15 | 0.92 | 5.64 | 2.27 | SFHFKEAWKHAIQKAKHMPDPWAEFHLE/KEEEIEEMEEEFEELIEERLKTLPPEAAEKVREIIERAKERLEKNPLTPEFVKELEEEVREEFLALHEALAAKLGKAVTPEAKKLIEEWVKLEIEMMTVIKLAKL |
| 16 | 0.89 | 23.86 | 34.67 | SFHFKEAWKHAIQKAKHMPDPWAEFHLE/GLAAVTAAALAELKALAKTPLQKAFLAAAERLVANPDATREEVIALFEELREKALELGYELLAEDVEEFIEELEERADLPRELLRRRALRKLLELAYKEQEIEKA |
| 17 | 0.81 | 19.51 | 3.93 | SFHFKEAWKHAIQKAKHMPDPWAEFHLE/GLEEKEKELLEKLKKLSKTPLRKKLLEVAEKIVKNKDATLEEILELLEELEKFFKELGDELLAEEVKKFKEELKKRKSLSKLVLRRRTLVFLLNLSLADNEIEEA |
| 18 | 0.92 | 22.17 | 31.10 | SFHFKEAWKHAIQKAKHMPDPWAEFHLE/GLEEKREEALARLEALARTPSQRKLLEAARRIVENEDATREEVIELLEEVREFFLEEGNELLAEEVEEFIEEVKKESSLSKLLLRRRTLVFLLELAEAEAEIEEA |
| 19 | 0.85 | 24.11 | 16.68 | SFHFKEAWKHAIQKAKHMPDPWAEFHLE/GLEEVEEEELKRLRELAKTELRKKFLEAAERIVENPEATREEIIELLEEVREKALELGNELLAEDVEKFIEEVRKRKDLSKLVLRRRALKFLLELVEAENEIEEA |
| 20 | 0.85 | 16.01 | 19.67 | SFHFKEAWKHAIQKAKHMPDPWAEFHLE/GLEAVEAAELAELEALAKTPLRKALLAVARTVVENPDATREELIALAEKLREKFKELGDFLLAEEVEKFIEELKEQSSLSPLVMRRRLLRFLRELALADNEIEKA |
| 21 | 0.82 | 23.61 | 28.53 | SFHFKEAWKHAIQKAKHMPDPWAEFHLE/GLEAKTAQLLAELEAKAKTPLQKALLEAARTVVANPEATLEELLAIVEE |

| | | | | |
|---|---|---|---|---|
| | | | | LEKKALELGDELLAEDLREFAKELRERSSLSPLLLRRRFL KFLLELADKVNEIEEA |
| 22 | 0.87 | 14.92 | 9.83 | SFHFKEAWKHAIQKAKHMPDPWAEFHLE/GLAAVEAAE LKRLKAMAKTPLRKALLAAAEEVLANPEATREELIALLK ELQKKALELGNELLAEEIKEFIKELEERADDSPLLLRRRV LKFLLELAEAEAEIEEA |
| 23 | 0.76 | 23.13 | 8.63 | SFHFKEAWKHAIQKAKHMPDPWAEFHLE/GLAAKTAAA LAALKAAAATPARKALLAAAQRVVAAPDATRDELLALL EELRAFFLAEGNELLAEEVEEFIAELEARAGLSKELMRRL TLRRLGELAAADAEIAAA |
| 24 | 0.88 | 19.49 | 25.09 | SFHFKEAWKHAIQKAKHMPDPWAEFHLE/AAAAAAAAE RAALLAAVRATAKAAREAILKLKDEEEAIKKILELFKAFL AALGREELAEDVERLAKELVEEGFSVEQIFRALFYYLMT RLGFSEELAEAILAEVE |
| 25 | 0.92 | 14.91 | 5.20 | SFHFKEAWKHAIQKAKHMPDPWAEFHLE/GLAAALAAE RAALRARLEAVLRAALEAILELEDEEEAIERIIELFTAFLD ELGDEEGKEYIKELAEELIKEGFSVYQILRALFIAVAELLG FSEEETEEMLERAL |
| 26 | 0.83 | 14.74 | 6.55 | SFHFKEAWKHAIQKAKHMPDPWAEFHLE/MEEELEEEEE EELLELIKEVLKKAREIIEKEKDEEKAIEKIVELFLEYFKKL GLEEYAKEVKKLAEELKKEGFSVWQILRALFIYTGTLLGI SEEKTEELIEEAE |
| 27 | 0.82 | 15.13 | 10.26 | SFHFKEAWKHAIQKAKHMPDPWAEFHLE/MAAALAAEK RAALLEKLRKTLKKVLKAIKELKDEDEAIKKILELFQKFF KELGDEEGAEYIKKLVEEMIKEGFSVVQIARAVFIAVGEL LGISEEETEALIKEAL |
| 28 | 0.82 | 14.44 | 5.59 | SFHFKEAWKHAIQKAKHMPDPWAEFHLE/GLAALLAAE KAALLARLEAVVKEMRKAILEEKDREEAIKKIEELFLKFT EELGLEEFGEYIKKLIEEMIEEGFSVEQILRATFIALAELLG ISEEETEAILERVL |
| 29 | 0.89 | 12.92 | 5.77 | SFHFKEAWKHAIQKAKHMPDPWAEFHLE/MKLLEEKKK KEELLKKLEEVVKKVLKAILELEDEEEAVAKILELFEAYL KELGLEEGAENVRKLAEEMIKEGFSVWQIARALFIYVSEL LGIDEETTEEILKKAE |
| 30 | 0.89 | 14.87 | 3.98 | SFHFKEAWKHAIQKAKHMPDPWAEFHLE/MEKEKEEKK KKELLEKIEETLKKARKAIKELEDEEEAIKEILKLFEEFFK ELGLEEFQKYIKKLIKELIEEGFSVEQILNAVFYAAMELLG ISEEEAEKIIEKVY |
| 31 | 0.85 | 13.02 | 6.24 | SFHFKEAWKHAIQKAKHMPDPWAEFHLE/MSHLLEELKR KELLERVKKVVKEVVKAILELEDEDEAIEKILKLFLEYMK ELGLEEHAEYIKRLAEEMKKEGFSVWQIAHALFIATMTL LGIDEETTEKILEEAL |
| 32 | 0.66 | 21.42 | 7.46 | SFHFKEAWKHAIQKAKHMPDPWAEFHLE/SLLEELEKLK AEHAEEWKLLEEYLKEVEEKEEELDKESILELFFEKAEPY |

| | | | | |
|---|---|---|---|---|
| | | | | LKKDPLKTVTNLAKVASAELEYLEKFAPLSEWKKKRILY TIKEALEEAKEELEKK |
| 33 | 0.71 | 17.60 | 3.64 | SFHFKEAWKHAIQKAKHMPDPWAEFHLE/SLLEEYEELK EEHEEEWELVEEYLEEVEEKWDELDKESILELFVKKLEPF LKKDPEETLKKIAEVLAAEAKLLEEFADLSSWKTLRILLT LRDFAEEALEELKKK |
| 34 | 0.63 | 20.35 | 10.17 | SFHFKEAWKHAIQKAKHMPDPWAEFHLE/DLAAALAAL RAAHAAEAALLDEYLAEVAARREELTLEQKLELFWEKL EPALKADPRATLAALAAVAEAEAAALRATAPLAAAPTL RDLETFAAALREALAALEAA |
| 35 | 0.67 | 18.12 | 10.36 | SFHFKEAWKHAIQKAKHMPDPWAEFHLE/SLEKELEELK KKLAEHFALLEEYLKLVEENKEELDLESKLELFWKLLEP YLKKDPLKTLENVHKVLEAELKYIEEKKPLSYAPKQRILR TFAAATKEAIEELKKK |
| 36 | 0.74 | 14.03 | 1.97 | SFHFKEAWKHAIQKAKHMPDPWAEFHLE/ELLEELEKLE EKYKEYVELLEEYLELVEENREELDLESKLELLFKLLEPF LKKDPLETVEKLALVAEAKARYLETFAPLSSWPLLRDLR TFNAALKEAGKELKKK |
| 37 | 0.79 | 13.22 | 2.87 | SFHFKEAWKHAIQKAKHMPDPWAEFHLE/SLLEELKKLK EEHKEEWKLLEEYLKYVEENKEELDKESKLELLFKKLEP YLKKDPVKTLKNLAKVAKAEAEYLKKFAPLSAAPKLRD LETFKAVLEEALKELEKR |
| 38 | 0.78 | 12.42 | 5.65 | SFHFKEAWKHAIQKAKHMPDPWAEFHLE/SLLEELEKLR KEHEEYVKLVEEILEEIKENKEKLDKESILELAFEKLEPYL KENPIETLENLAKVLKAELEYLEKFAGLEAWPKQRILYTL REAVEEALEELKKR |
| 39 | 0.59 | 19.51 | 16.39 | SFHFKEAWKHAIQKAKHMPDPWAEFHLE/SLEEELEKLK EELEEEYELLEEYLEEVKERWEELDKESILELFFKKLEPYL KEDPLKTLENLAKVAEAELEYLKKFAPLSKWPLARILETI RAALKEALKALKER |
| 40 | 0.88 | 19.37 | 12.16 | SFHFKEAWKHAIQKAKHMPDPWAEFHLE/SAEEERKLAL VEELAEKLEELEKLVDTAARDWELMERRGAPAALRAEL WAAFRERAGALLAEVEALLAELRALLPPEEAAEPLARLE ARLARVRARIAALEARAA |
| 41 | 0.84 | 20.05 | 14.40 | SFHFKEAWKHAIQKAKHMPDPWAEFHLE/SEREKRLKEL VEKTAEKLEELEKLVRTFLRHWDLLERRGAPAALRAELR ARFLATSSALLAEVKKLLEKLKKEAPEEEAKEFLARLEK RLAEVEALYAEALARWA |
| 42 | 0.88 | 10.01 | 6.08 | SFHFKEAWKHAIQKAKHMPDPWAEFHLE/SEREKRLKEL VEKLAEKLEELEKLVKTFLKNWELMEKKGAPQELIEKL WKEFKETSEKLLKEVKELLEKLKKEISPEEAKEFLKELEK RLKENEELIKELKERAK |
| 43 | 0.85 | 17.92 | 14.79 | SFHFKEAWKHAIQKAKHMPDPWAEFHLE/SEREERLKKL VEELWEKLEELNKLVETALRNLELMEKKGAPEELKEKL |

| | | | | |
|---|---|---|---|---|
| | | | | WKEFEERSTKLLKEVEELLKKLKEEISKEEAKEFLKEAEK KLEEVKKKIKELKERRA |
| 44 | 0.85 | 18.67 | 13.46 | SFHFKEAWKHAIQKAKHMPDPWAEFHLE/SSREERLLAL VEETAEALEELEKLVRLFRRHLALMARRGAPQALRDQL WADFRARASALLARVKALLEELRKEAPPELAKEFLARLE ERLAEVEALIAELEAAAA |
| 45 | 0.86 | 19.39 | 34.41 | SFHFKEAWKHAIQKAKHMPDPWAEFHLE/SARREEWLEL VEKTAEALEELKKLVDTYLRNLDLMEERGAPQALRDRL RAEFEATSSALLKEVKKLLEELKKKLPKEEAKEFLKRLEE ALKEVEAKIAELRARFA |
| 46 | 0.87 | 16.13 | 10.55 | SFHFKEAWKHAIQKAKHMPDPWAEFHLE/SSREERLLRL VEELAEALEELQKLVETAARNLALMAARGAPQALIDAL WADFRERAGALLAEVEALLARLRAEAPAEELAEPLAELE ARLAEVKALIAALEAAAA |
| 47 | 0.87 | 22.70 | 8.71 | SFHFKEAWKHAIQKAKHMPDPWAEFHLE/SEREKELKEL VEKTAEALEELSKLVDLALRHWELMERRGAPEELRREL WARFRATSTALRAEVKALLKELKEKISPEEGKEFLKRLE ARLKENEAKIAELEERAA |
| 48 | 0.75 | 12.61 | 5.22 | SFHFKEAWKHAIQKAKHMPDPWAEFHLE/KELEELKKLV EIVKKMLENEKNEELFEETLKIMKEISEELAELVKLIIEAIK ALLEGDKEKAIEYIDKAYELVLKSKDRAATALYHEFMKY LEKKYKELEEEEKK |
| 49 | 0.90 | 8.32 | 5.31 | SFHFKEAWKHAIQKAKHMPDPWAEFHLE/SELEKLLELV EKVKEALENENDEEKWKELIEYLKEISEELAELAEIIVNA VKAYLEGNKEEMIEYIDKADELVKKLGSRAATALLHEFY DWLEKKLKELEEEEKK |
| 50 | 0.81 | 9.55 | 2.05 | SFHFKEAWKHAIQKAKHMPDPWAEFHLE/SELARLAELV ALVEEALEAAEDEALHAAVLARFAAISPELAELAKLIVK AVKALLEGKKEEAIKYIDEAYELVQKSGDVAARAVFLQF MAWLEKKYKELEEEEEK |
| 51 | 0.90 | 8.69 | 3.51 | SFHFKEAWKHAIQKAKHMPDPWAEFHLE/SELEEDLRLV ERFKEALENEEDEELHEEFLEEMLEISEELAELVEIAIKAIK ALNEGDVEKFIKYIDELYKLVEELGDRRALALAHEFFEW LEKKYKELKEKEEK |
| 52 | 0.90 | 9.31 | 3.27 | SFHFKEAWKHAIQKAKHMPDPWAEFHLE/SSIEKYLEAIE LVREALKNEKNEELQKKVIEYFKKISEELAKLIEIIFKAIK AINEGKKEEFIKYIDEAYELVLKSGDKLMLAVFHEFMEW LEEKYKKLEEEEKK |
| 53 | 0.90 | 9.26 | 3.69 | SFHFKEAWKHAIQKAKHMPDPWAEFHLE/SALEEDLELV ELMREALEKEEDEEFHKEVLEKMAEISEELAELVKIIIEAI KALLEGNKEKFIEYIDKAYELVQELKSRLAQALYHEFME WLEKKYKELEEKEKA |
| 54 | 0.90 | 9.79 | 3.00 | SFHFKEAWKHAIQKAKHMPDPWAEFHLE/AELEEYKEIIE KMKEALKNEEDEELWEEVIKYMKEKSETLAELVELIKEA |

| | | | | |
|---|---|---|---|---|
| | | | | IKALLEGKKEEFIEYIDKAYELVEKDKDRLSLALYLEFMK WLEERYEKELKKEKE |
| 55 | 0.90 | 22.09 | 10.82 | SFHFKEAWKHAIQKAKHMPDPWAEFHLE/EELELYEKLV ALFKEALENEENEELKKALLEEFKKISEELAKLVELIFKAI EAYLEGKKEEMIKYIDEAYKLVLELGDRLALLLFLEFME WLKKKYEELEKKAKK |
| 56 | 0.87 | 12.07 | 3.66 | SFHFKEAWKHAIQKAKHMPDPWAEFHLE/KKTEAAKEL KALLDKLKELIKNRIKLYEEGESIEEKLDEIKKELIEALKK LGVPEEAIKEVEKLFEELKKHIKKIIEGEKEENADKAVEA LKKLEEAVKAAIEAL |
| 57 | 0.89 | 9.89 | 3.80 | SFHFKEAWKHAIQKAKHMPDPWAEFHLE/KKKEIAKKV KAKLEEIKELIFNTIELVEKGEDIVKKFKELKEKLIKLLKEI GFPEEVIKEVKELFDKMIEAIEKIIEGEKEENAEKLVEYFE KLEEKLKEAVEEL |
| 58 | 0.88 | 8.68 | 3.64 | SFHFKEAWKHAIQKAKHMPDPWAEFHLE/KKREIAKEIR EKIDEIKELLENRIELIEKGEDVEKKFEELKKELIEKLKEA GFPEEVIKEVKELFDKLIEAIKKIEEGEKEENAEEAVEYLK KLEEKLKEAVEKL |
| 59 | 0.88 | 8.75 | 2.45 | SFHFKEAWKHAIQKAKHMPDPWAEFHLE/KLREQIKELK KLLDKLKELIENSIELYEKGENIEEEFEKLKKELLKKLKEL GVPEEEQKKVEELFDKMIEAIKKIIEGDKEENADELVKNF KELEEALKEAAKKL |
| 60 | 0.87 | 7.82 | 2.74 | SFHFKEAWKHAIQKAKHMPDPWAEFHLE/AARAALREL RALLDELRELLENSIKLVEEGKDIRAELEALRARLLAALA ALGVPAADRARVAALFDRLLEHIERIIAGEKEESADEAVE ALAALEAALAEAARAL |
| 61 | 0.86 | 8.67 | 2.01 | SFHFKEAWKHAIQKAKHMPDPWAEFHLE/ELRKIAEELK ALIEKLKELLENRLKLVEEGKDVEAELDALKKELLAKLK EAGFPEEKREKAKALFDEMKEAIKKIIEGEKEENADKLVE SLKELEEELLKEGVKEL |
| 62 | 0.87 | 10.13 | 3.07 | SFHFKEAWKHAIQKAKHMPDPWAEFHLE/SLREIIKELKE KIDELKELIENRIKLKEEGEDVEKKLEELKKKLIELLKKA GFPEEVIKKVEALFEELIKHIKKIEEGEKEENADKAVEAL KALEETLKEAAKAL |
| 63 | 0.87 | 9.15 | 3.14 | SFHFKEAWKHAIQKAKHMPDPWAEFHLE/KLREIAIEVK KKLDELLELIEKRIELVEKGESVEEELKKLKEELLKLLEK AGFPEEEREKVEKLFNELIEHLKKIEEGEKEENADKAVEA LKELREALKEAVESL |

*CAM1/2 scores and sequences shaded.

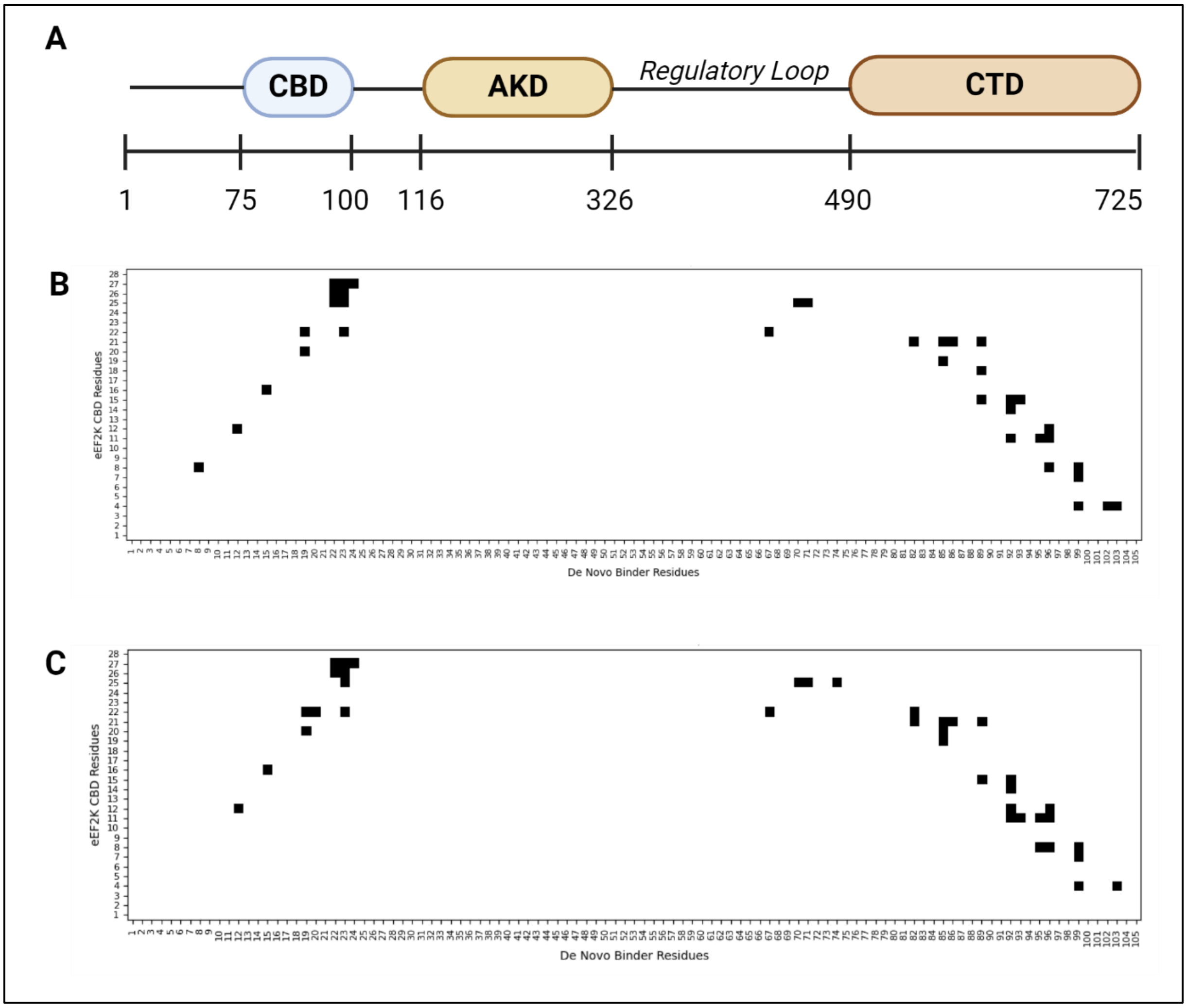


**Figure S1. Predicted contacts of eEF2K CBD with CAM1/2.** A) Domain map of the alpha kinase eEF2K, where CBD, AKD, and CTD, represent the calmodulin binding domain, alpha kinase domain, and c-terminal eEF2 binding domain, respectively. B) Protein-protein contact map between CBD residues and CAM1 or C) CAM2. Contacts displayed are less than or equal to 8 (Å) between backbone alpha-carbons.

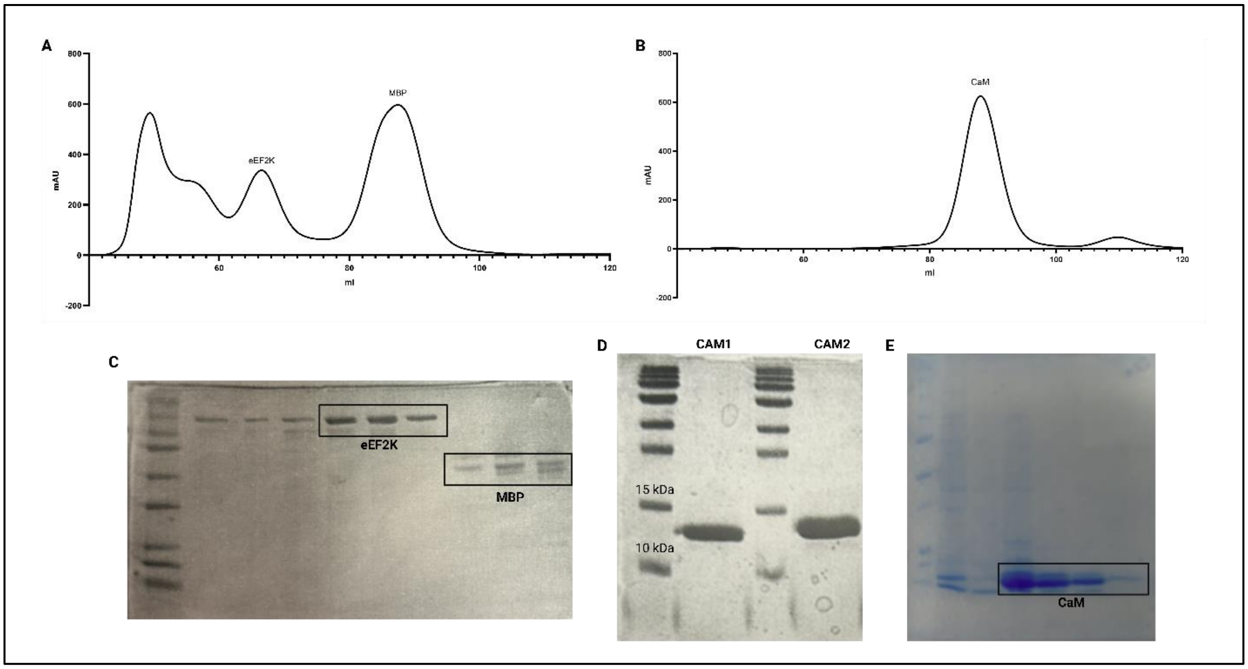


**Figure S2. Recombinant protein production of eEF2K, CAM1/2, and CaM.** A) Size-exclusion chromatogram of eEF2K and MBP after cleavage of MBP with TEV protease and B) calmodulin. C) Purified fractions of eEF2K, D) CAM1 or CAM2, and E) calmodulin (purified with nickel chromatography then size-exclusion).

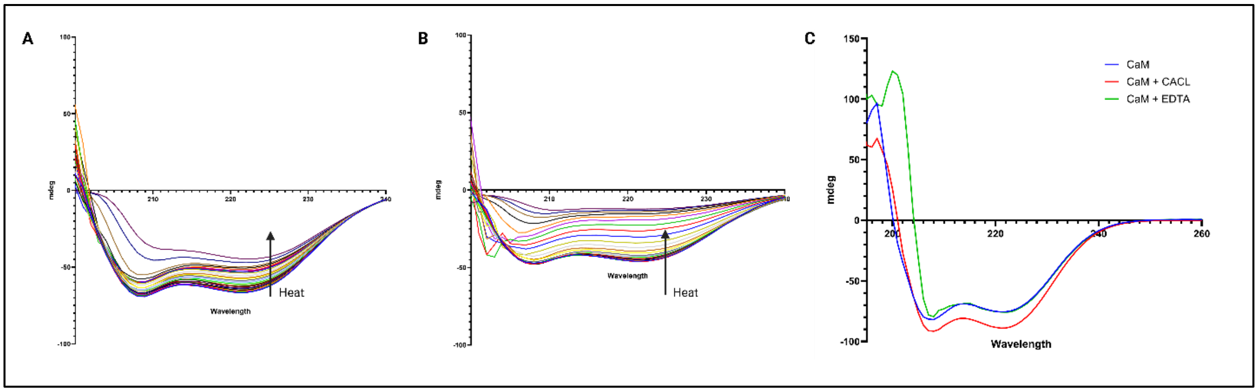


**Figure S3. Circular dichroism spectroscopy studies of CAM1 and CaM.** Denaturation circular dichroism (CD) spectra's of A) CAM1 and B) calmodulin. C) CD spectra of calmodulin treated with 1 mM CaCl or 1 mM EDTA.

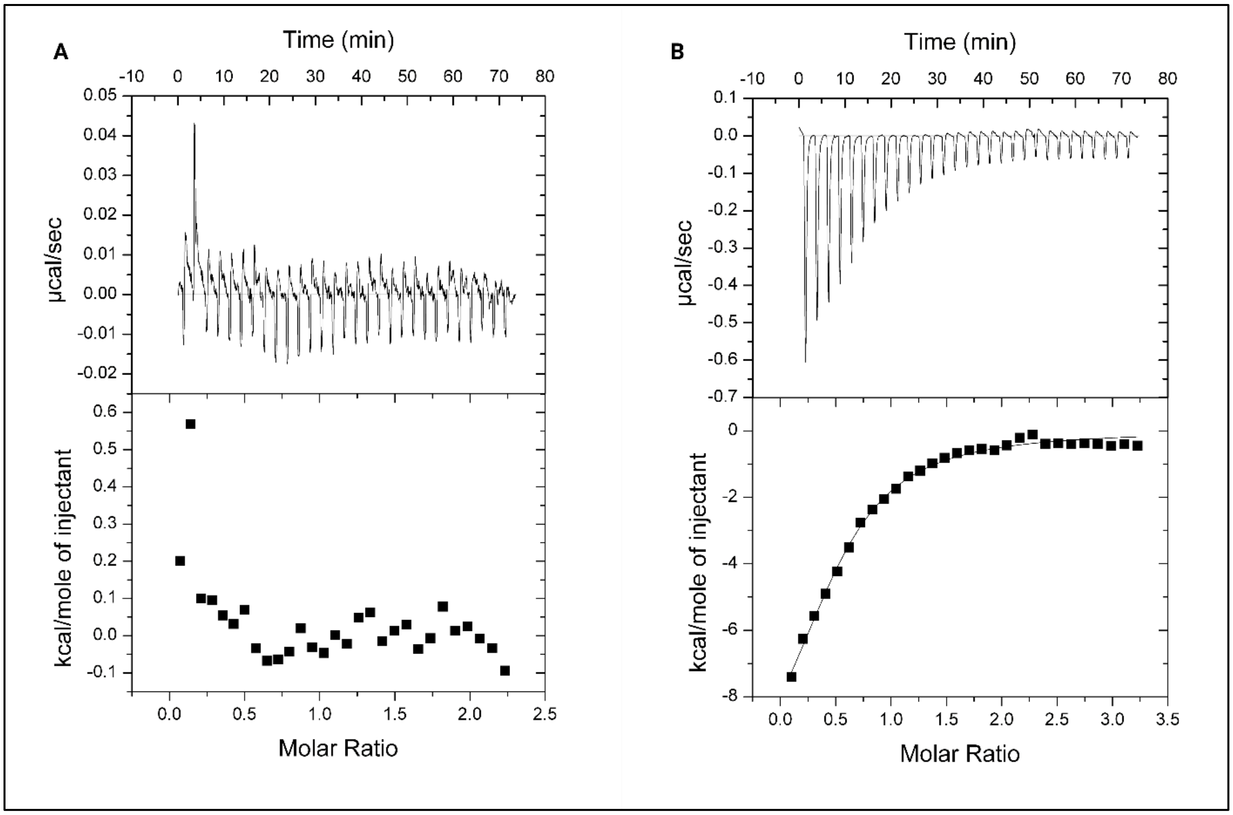


**Figure S4. Isothermal calorimetry data.** A) No binding signature detected between free calcium and CAM1. B) Calmodulin binds to the CBD of eEF2K.
